# Identification of bioactive metabolites in human iPSC-derived dopaminergic neurons with PARK2 mutation: altered mitochondrial and energy metabolism

**DOI:** 10.1101/2020.07.10.151902

**Authors:** Justyna Okarmus, Jesper F. Havelund, Matias Ryding, Sissel I. Schmidt, Helle Bogetofte, Nils J. Færgeman, Poul Hyttel, Morten Meyer

## Abstract

*PARK2* (parkin) mutations cause early onset of autosomal recessively inherited Parkinson’s disease (PD). Parkin is an ubiquitin E3 ligase and has been reported to participate in several cellular functions, including mitochondrial homeostasis. However, the specific metabolomic changes caused by parkin depletion remain largely unknown. Human induced pluripotent stem cells (iPSCs) with *PARK2* knockout (KO) provide a valuable model for studying parkin dysfunction in dopaminergic neurons. In the current study, we used isogenic iPSCs to investigate the effect of parkin loss-of-function by comparative metabolomics analysis. The metabolomic profile of the *PARK2* KO neurons differed substantially from that of healthy controls. We found increased tricarboxylic acid (TCA) cycle activity, perturbed mitochondrial ultrastructure connected with ATP depletion, glycolysis dysregulation with lactate accumulation, and elevated levels of short- and long-chain carnitines. These mitochondrial and energy perturbations in the *PARK2* KO neurons were combined with increased levels of oxidative stress and a decreased anti-oxidative response. In conclusion, our data describe a unique metabolomic profile associated with parkin dysfunction, demonstrating several PD-related cellular defects. Our findings support and expand previously described PD phenotypic features and show that combining metabolomic analysis with an iPSC-derived dopaminergic neuronal model of PD is a valuable approach to obtain novel insight into the disease pathogenesis.

## Introduction

The pathological hallmark of Parkinson’s disease (PD) is the marked loss of dopaminergic neurons in the substantia nigra pars compacta (SNc), causing dopamine depletion in the striatum (1–4). Although the multifactorial etiology of PD and the pathological mechanisms underlying the neuronal degeneration remain largely undetermined, various PD-related genetic-environmental interactions are thought to contribute to the pathogenesis of the disease. Mitochondrial dysfunction associated with oxidative stress and energy failure is increasingly thought to be implicated in PD (5–9).

Mutations in a number of genes have been found to cause monogenic forms of PD with both recessive (e.g. *PARK2, PINK1, DJ-1*) and autosomal dominant transmission (e.g. *LRRK2, SNCA*) (10, 11). Loss-of-function mutations of the *PARK2* gene, encoding the cytosolic E3 ubiquitin protein ligase parkin, account for a large proportion of familial early-onset PD cases. Even though monogenic (often familial) forms of PD are not frequent (less than 5% of all PD cases), elucidation of their molecular mechanisms could help identify causes of the more common sporadic forms of the disease (12, 13). The protein parkin is thought to play an essential role in the regulation of mitochondrial homeostasis and dynamics. Increased expression of the *PARK2* gene transcript confers protection from stress-induced cell death (5, 6, 8). Furthermore, loss of parkin increases susceptibility to stress and death in primary cells (9, 14, 15). It is thus believed that *PARK2*-genetic causes of PD likely involve a loss-of-function phenotype that leads to the clinical presentation of the disease (3, 16).

Mitochondria are the powerhouses of the cell and play an important role in various metabolic pathways. Apart from their crucial role in cellular energy metabolism (ATP production via oxidative phosphorylation (OXPHOS)), mitochondria are powerful organelles that regulate reactive oxygen species (ROS) production, NAD+/NADH ratio and programmed cell death (14–16). Disturbances in mitochondrial function have been suggested to contribute to neurodegeneration (17). Evidence of mitochondrial impairment in PD comes from mitochondrial toxin-induced models of the disease (18, 19), and from the examination of mitochondria in post-mortem tissue from patients with idiopathic PD (20, 21). Disease-causing mutations in genes encoding for proteins with a mitochondrial function (*PARK2, PINK1, DJ-1, HTRA2*) have also been identified, further supporting the conclusion that mitochondrial dysfunction is a hallmark of PD (22–25). Our recent studies of induced pluripotent stem cell (iPSC)-derived dopaminergic neurons with *PARK2* knockout (KO) have reported abnormal mitochondrial morphology and function associated with increased oxidative stress and autophagic-lysosmal perturbations (26, 27). In mammalian cells it has been shown that dissipation of mitochondrial membrane potential (MMP) leads to recruitment of parkin to mitochondria, followed by their elimination by selective autophagy (mitophagy) (28). Imapirement of mechanisms involved in removal of damaged mitochondria may lead to increased generation of ROS and irrevocable apoptotic cell death, which has been reported for dopaminergic neurons in PD (29).

The emergence of metabolomic technologies enables systematic measurement of low-molecular-weight compounds to provide an overview of alterations in metabolic pathways induced by a given perturbation, which could be a gene mutation or infliction of a sporadic disease condition (30). Metabolomics is therefore useful for studying neurodegenerative disorders as it can reveal disease pathomechanisms and identify potential biomarkers (31, 32). Studies of cerebrospinal fluid (CSF) from patients with PD have identified PD-specific alterations in several metabolic pathways, including polyamine metabolism, the purine, pyruvate, kynurenine pathways, and redox markers (31, 33–36). Although there is growing interest in the use of metabolomics in PD research, many of these studies do not corroborate each other possibly due to low sample numbers, clinical heterogeneity, and different analytical approaches (30).

To our knowledge, no previous studies exist on the metabolomic profile of human iPSC-derived neurons with parkin deficiency. In the present study, we have aimed at demonstrating disease-relevant cellular effects of parkin dysfunction in dopaminergic neurons. To improve our understanding of the parkin-mediated PD phenotype, we describe here the metabolic profile of a *PARK2* KO iPSC line compared to its healthy control. We report a metabolic signature consisting of 92 compounds of which 51 metabolites significantly distinguish the *PARK2* KO neurons from healthy controls. Our study identifies systemic, metabolic pathway derangements that are possibly caused by primary mitochondrial changes and suggests how disturbances in mitochondrial homeostasis contribute to PD. These findings represent a significant contribution to our understanding of PD pathogenesis and also suggest that PD could be considered a metabolic disorder.

## Results

### Metabolomic profiling of *PARK2* KO neurons by LC-MS

To gain new insight into the role of parkin dysfunction in PD, we applied liquid chromatography-mass spectrometry (LC-MS) metabolomics analysis to compare *PARK2* KO neurons with healthy controls. An overview of the LC-MS metabolomics workflow is presented in *Fig. 1*. Neurons were generated using a previously described robust differentiation protocol (26, 37). Differentiated cells were collected, and the metabolites were extracted for LC-MS analysis. Of the 92 structurally annotated metabolites, 51 compounds significantly differentiated *PARK2* KO neurons from controls (q-value < 0.05) (*Supp. Table 1*). Principal component analysis (PCA) was used to investigate general interrelationships between the groups, including clustering and outliers among the samples. PCA revealed a clear separation of metabolomes between *PARK2* KO neurons and control neurons (*Fig. 2A*). A clustered analysis heatmap was generated to group related compounds and illustrate the differential profiles (*Fig. 2B*). Following hierarchical cluster analysis, Metabolite Set Enrichment Analysis (MSEA) was performed on detected metabolites to identify which pathways were affected by the distinguishing metabolites (*Fig. 3A*). In parallel, we also utilized the Metabolomic Pathway Analysis (MetPA) module of MetaboAnalyst that combines results from the pathway enrichment analysis with the pathway topology analysis. A graphical list of the pathways identified and their relative impact is shown in *Fig. 3B*. The most important pathways (FDR < 0.05; impact values > 0.1) included: (i) tricarboxylic acid (TCA) cycle; (ii) lactate-pyruvate metabolism; (iii) glycolysis-gluconeogenesis metabolism; (iv) glutathione metabolism; (v) carnitine and acetyl groups; (vi) alanine, aspartate, and glutamate metabolism; (vii) glyoxylate and dicarboxylate metabolism. Taken together, these results demonstrate the capacity of LC-MS to identify a large number of metabolites and metabolic pathways implicated in PD, providing a valuable resource for further investigation.

**Figure 1:**
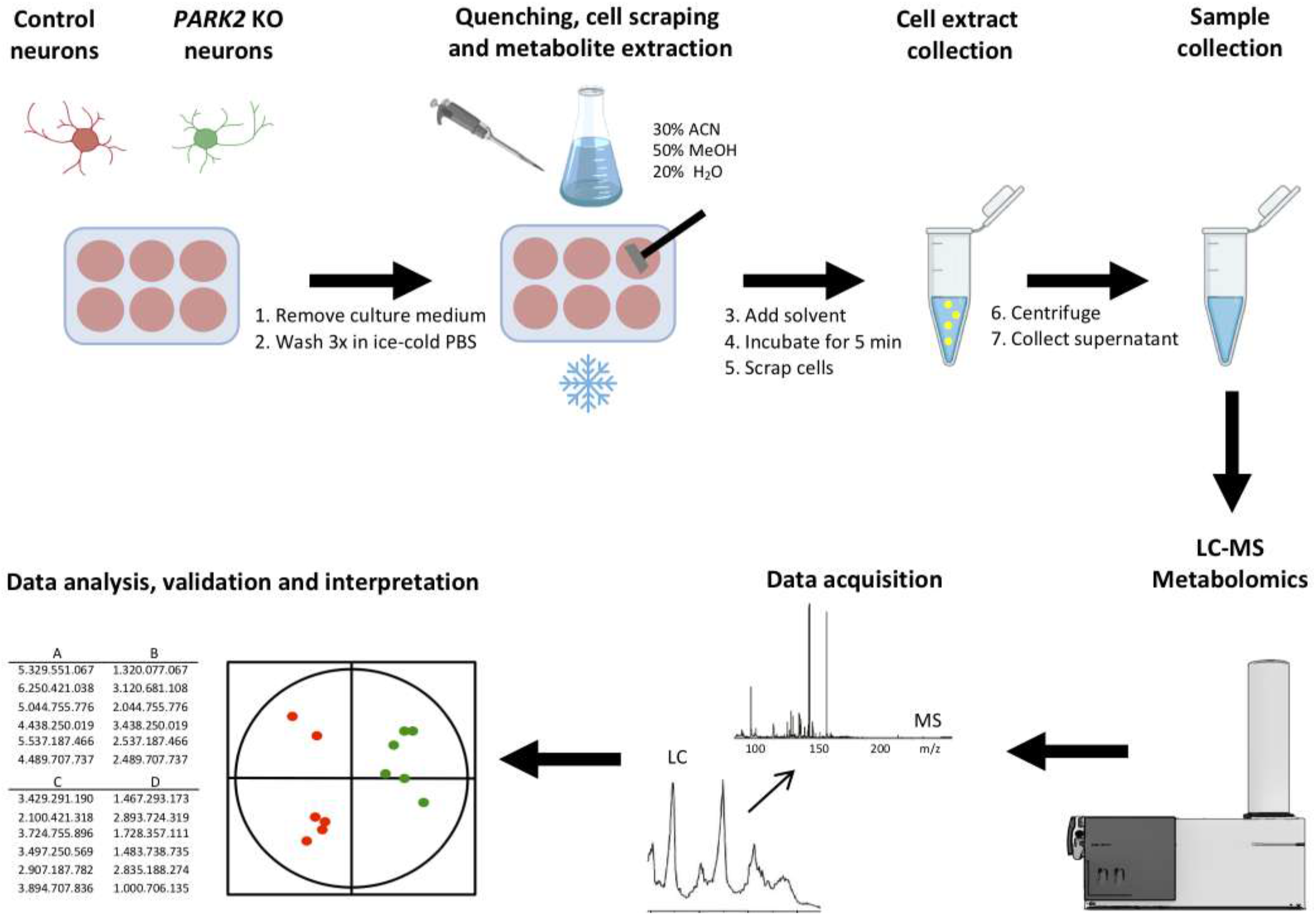
The workflow of liquid chromatography-mass spectrometry (LC-MS) metabolomics analysis.

**Figure 2:**
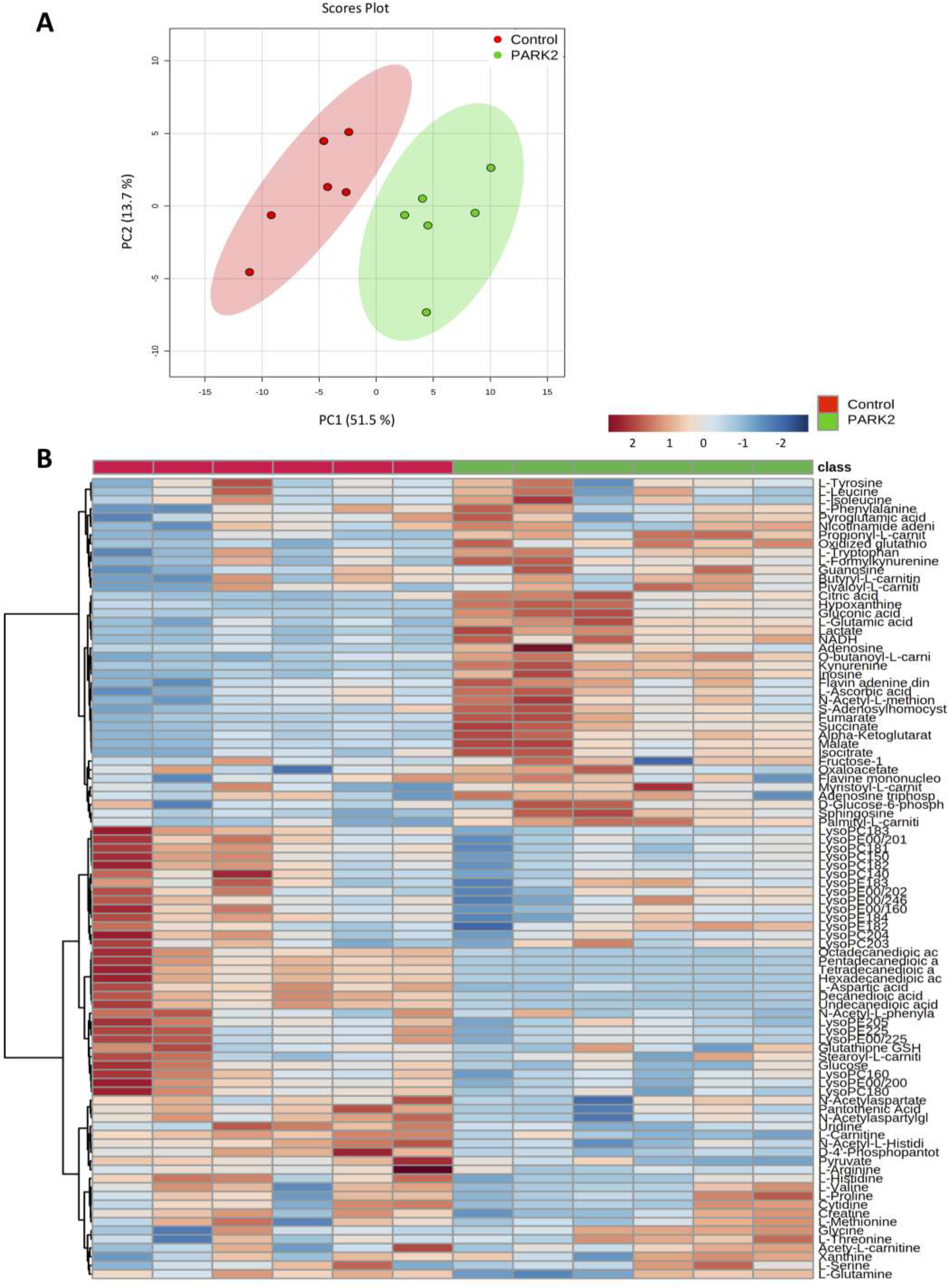
Comparative metabolic profiling of isogenic *PARK2* KO and control neurons. A) Principal component analysis (PCA) score plot showing a significant separation between *PARK2* KO neurons and control neurons. The observations are coded according to class membership: green = *PARK2* KO neurons, red = control neurons. B) Heat map of detected metabolomics dataset in the *PARK2* KO neurons and control neurons. The heat map depicts high (red) and low (blue) relative levels of metabolite variation. Individual compounds (vertical axis) are separated using hierarchical clustering (Ward’s algorithm) with the dendrogram being scaled to represent the distance between each branch (distance measure: Pearson’s correlation).

**Figure 3:**
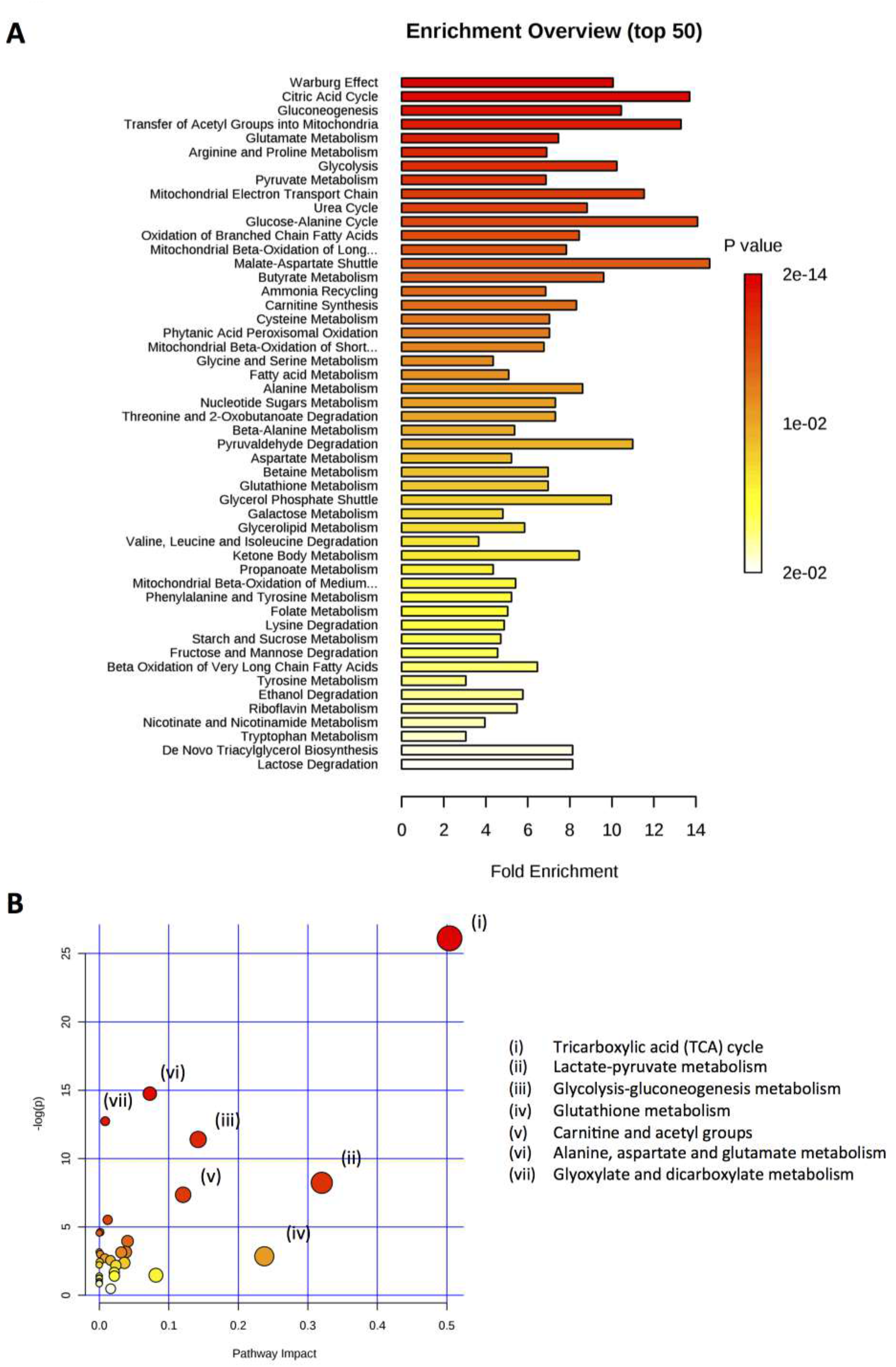
Metabolite associated pathways. A) Summary plot for the metabolite set enrichment analysis (MSEA) ranked by Holm p-value and showing top 50 perturbed pathways (generated by MetaboAnalyst software package). Color intensity (white-to-red) reflects increasing statistical significance. B) Metabolic pathway analysis (MetPA). All the matched pathways are displayed as circles. The color and size of each circle are based on p-value and pathway impact value, respectively. The most impacted pathways with high statistical significance scores are indicated as follows: (i) tricarboxylic acid (TCA) cycle; (ii) lactate-pyruvate metabolism; (iii) glycolysis-gluconeogenesis metabolism; (iv) glutathione metabolism; (v) carnitine and acetyl groups; (vi) alanine, aspartate, and glutamate metabolism; (vii) glyoxylate and dicarboxylate metabolism.

### Increased tricarboxylic acid (TCA) cycle activity in *PARK2* KO neurons

TCA, also known as Krebs cycle, forms a major metabolic hub and is involved in many disease states involving energy imbalance (38). Using LC-MS metabolomic analysis, we detected seven out of eight TCA cycle intermediates: citrate, isocitrate, α-ketoglutarate, succinate, fumarate, malate, and oxaloacetate. The levels of all the detected TCA cycle intermediates, except oxaloacetate, were significantly elevated in the *PARK2* KO neurons. The upregulation was presented as fold change compared to healthy control neurons (*Fig. 4A*), and relative abundance was also depicted (*Fig. 4B*). These findings suggest that parkin deficiency promotes the metabolic flux of the TCA cycle.

**Figure 4:**
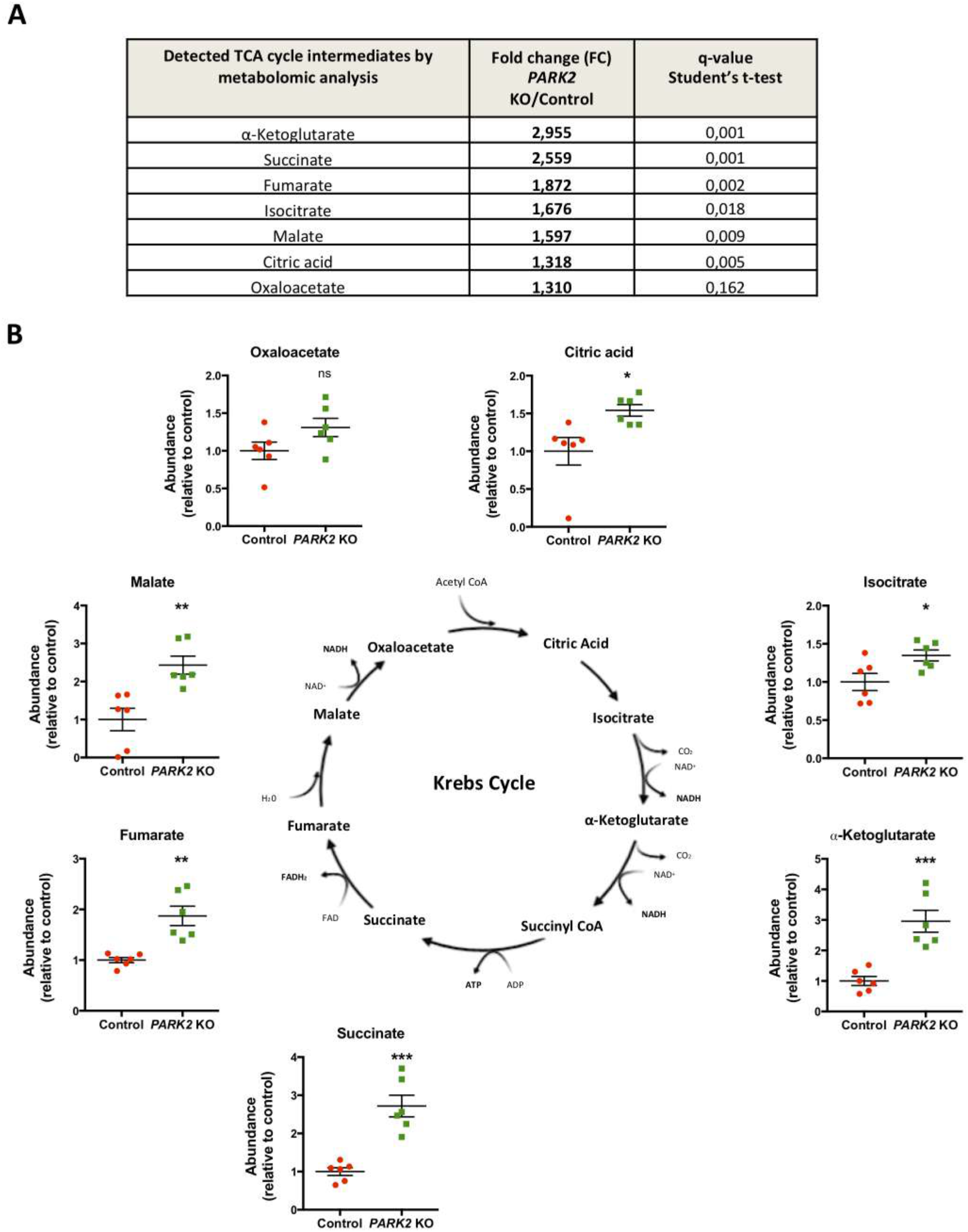
Metabolomic profile of the TCA cycle. A) Table listing fold change for detected TCA intermediates in metabolomics analysis. B) Graphical overview of TCA cycle indicating increased levels of its intermediates (citric acid, isocitrate, α-ketoglutarate, succinate, fumarate, malate, and oxaloacetate) in the *PARK2* KO neurons. Data presented as mean ± SEM, data from 3 independent differentiations, significant differences are indicated by *p < 0.05, **p < 0.01, ***p < 0.001, and ns: not significant, Student’s t-test.

### Mitochondrial membrane potential, mitochondrial ultrastructure, and ATP levels are perturbed in *PARK2* KO neurons

To study the role of parkin in mitochondrial maintenance and function, mitochondrial membrane potential (MMP) was monitored in *PARK2* KO and control neurons using tetramethylrhodamine methyl ester (TMRE), while FCCP treatment served as a negative control. When neurons were loaded with TMRE and examined by fluorescence microscopy, the *PARK2* KO neurons appeared to have diminished MMP (*Fig. 5A*). We verified this by a fluorescence plate reader-based quantification of the TMRE signal, which showed similarly reduced MMP (*Fig. 5B*). In addition to the functional mitochondrial analyses, comparative transmission electron microscopy (TEM) revealed clear changes in mitochondrial morphology in the *PARK2* KO neurons. Mitochondria in control neurons displayed a typical oval profile with well-organized cristae whereas mitochondria of *PARK2* KO neurons were swollen and exhibited aberrant cristae, disorganized inner membrane, and non-dense matrix (*Fig. 5C*). Based on these morphological observations, mitochondria were categorized into two groups: normal (with well-organized cristae) and abnormal (swollen with irregular cristae). Although quantification of mitochondrial number relative to cytoplasm showed no significant differences in total mitochondria per cell between *PARK2* KO and control neurons, there was a significant difference in the number of abnormal mitochondria (*Fig. 5D*). Since changes to the organization of the mitochondrial internal membrane are known to regulate mitochondrial respiratory function (39), the complexity of mitochondrial cristae was analyzed using the ratio between the mitochondrial cristae perimeter and the external perimeter. This ratio was significantly decreased in *PARK2* KO neurons compared to control (*Fig. 5E*), indicating loss of cristae complexity. Reduction in MMP and disruption of mitochondrial ultrastructure, including cristae complexity in the *PARK2* KO neurons, may suggest disturbances in the electron transport chain, which is crucial for ATP production (40). Therefore, to test energy production in the generated neurons, we measured the cellular ATP content. This was significantly decreased in neurons carrying *PARK2* KO as compared with control neurons (*Fig. 5F*), indicating an energy deficiency.

**Figure 5:**
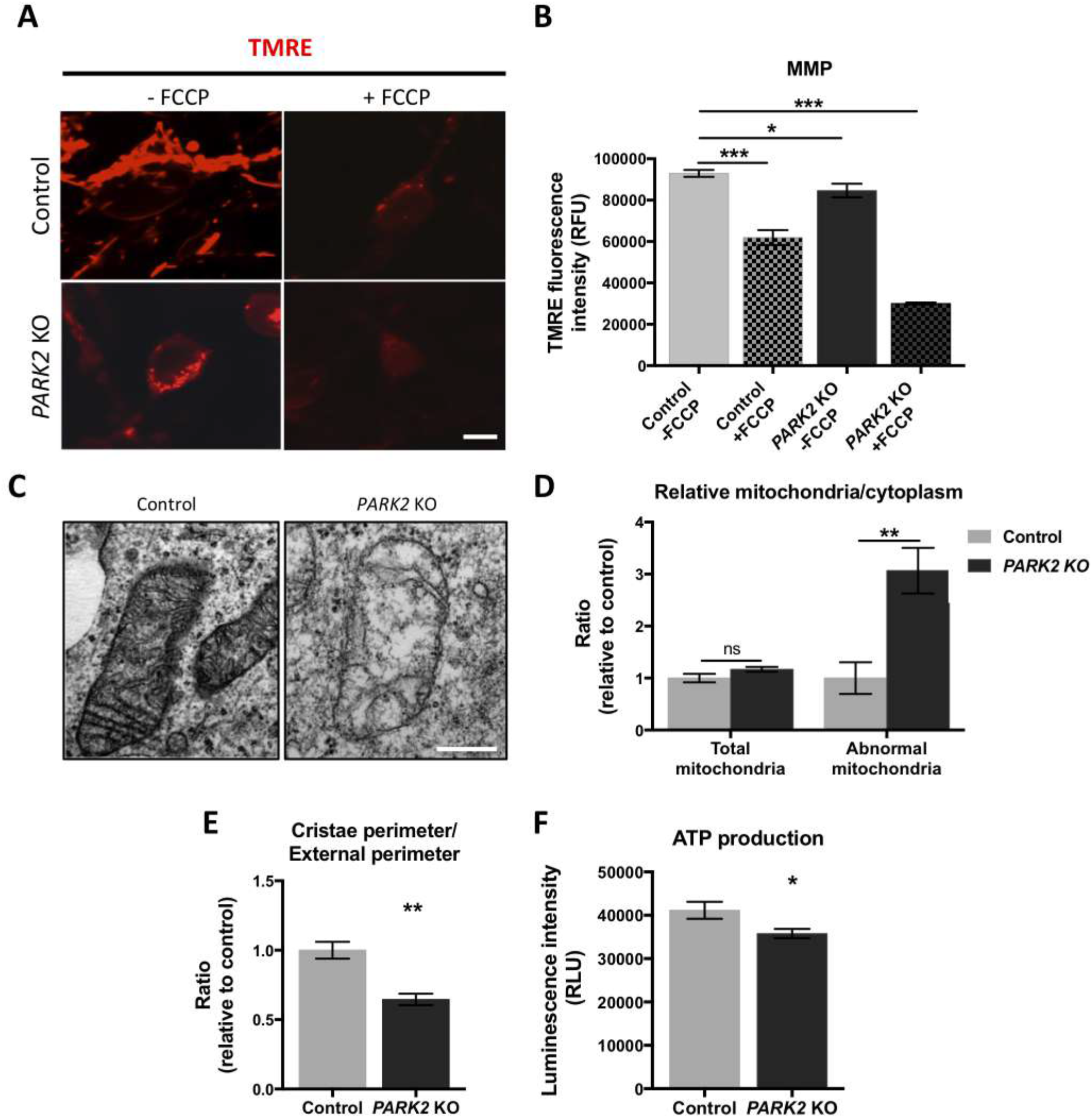
MMP, mitochondrial ultrastructure and cellular ATP levels are perturbed in *PARK2* KO neurons. A) Differentiated neurons were loaded with mitochondrial membrane indicator TMRE (50 nM). Changes in fluorescence were measured with (left panel) and without (right panel) application of the membrane potential uncoupler, FCCP (negative control). Scale bar: 10 μm. B) Graph presenting TMRE intensity from plate reader analysis. Data presented as mean ± SEM, data from 2 independent experiments, significant differences are indicated by *p < 0.05, ***p < 0.001, one-way ANOVA followed by Dunnett’s post hoc test for multiple comparisons. C) TEM pictures presenting mitochondria in control and *PARK2* KO neurons. Scale bar: 500 nm. D) Quantification of mitochondria/cytoplasm ratios showing the relative abundance of total and abnormal mitochondria in *PARK2* KO neurons. E) The ratio between the internal cristae perimeter and the external perimeter. F) Total ATP level determined using a luciferase assay kit in *PARK2* KO neurons. Data are presented as mean ± SEM, data from 3 independent differentiations, significant differences are indicated by *p < 0.05, **p < 0.01, Student’s t-test.

### Dysregulation of glucose metabolism and lactate accumulation in *PARK2* KO neurons

The detected TCA perturbations and ATP depletion correlated well with our earlier findings of decreased levels of a number of proteins of importance for energy metabolism (26). This prompted us to perform a joint protein–metabolite pathway analysis. Particularly, in the glycolytic pathway we found a high occurrence of both proteins and metabolites that were significantly dysregulated, including two important glycolytic enzymes, pyruvate dehydrogenase kinase (PKM) and lactate dehydrogenase (LDH), which were significantly reduced in the *PARK2* KO neurons (*Fig. 6*). When we compared the levels of glycolytic intermediates detected by the metabolomic analysis, we found that glucose levels were significantly decreased in the *PARK2* KO neuronal cultures (*Fig. 7A*). Further intermediates of the glycolysis pathway that were detected by LC-MS were D-glucose-6-phosphate, fructose-1,6-bisphosphate, and the end-product pyruvate. Although no difference in the level of D-glucose-6-phosphate was found (*Fig. 7B*), we detected a significant increase in the level of fructose-1,6-bisphosphate, an allosteric activator of glycolysis, in the *PARK2* KO neurons (*Fig. 7C*). Importantly, the abundance of pyruvate was reduced in the *PARK2* KO neurons alongside a highly elevated level of lactate. As lactate is a downstream by-product of pyruvate in the anaerobic glycolysis, this suggests perturbations in the glycolytic pathway and upregulation of lactate (*Figs. 7D, E*). To confirm our observations, we also determined lactate enzymatically. Consistent with our metabolomic measurements, we found that both intra- and extracellular lactate levels were significantly increased in the *PARK2* KO neuronal cultures compared to controls (*Fig. 7F*). Finally, we determined glucose uptake by measuring the uptake of the fluorescent glucose analogue 2-DG. No significant difference in glucose uptake was found between the two neuronal populations (*Fig. 7G*), indicating that glucose uptake was not responsible for driving the glycolytic changes observed for the *PARK2* KO neurons.

**Figure 6:**
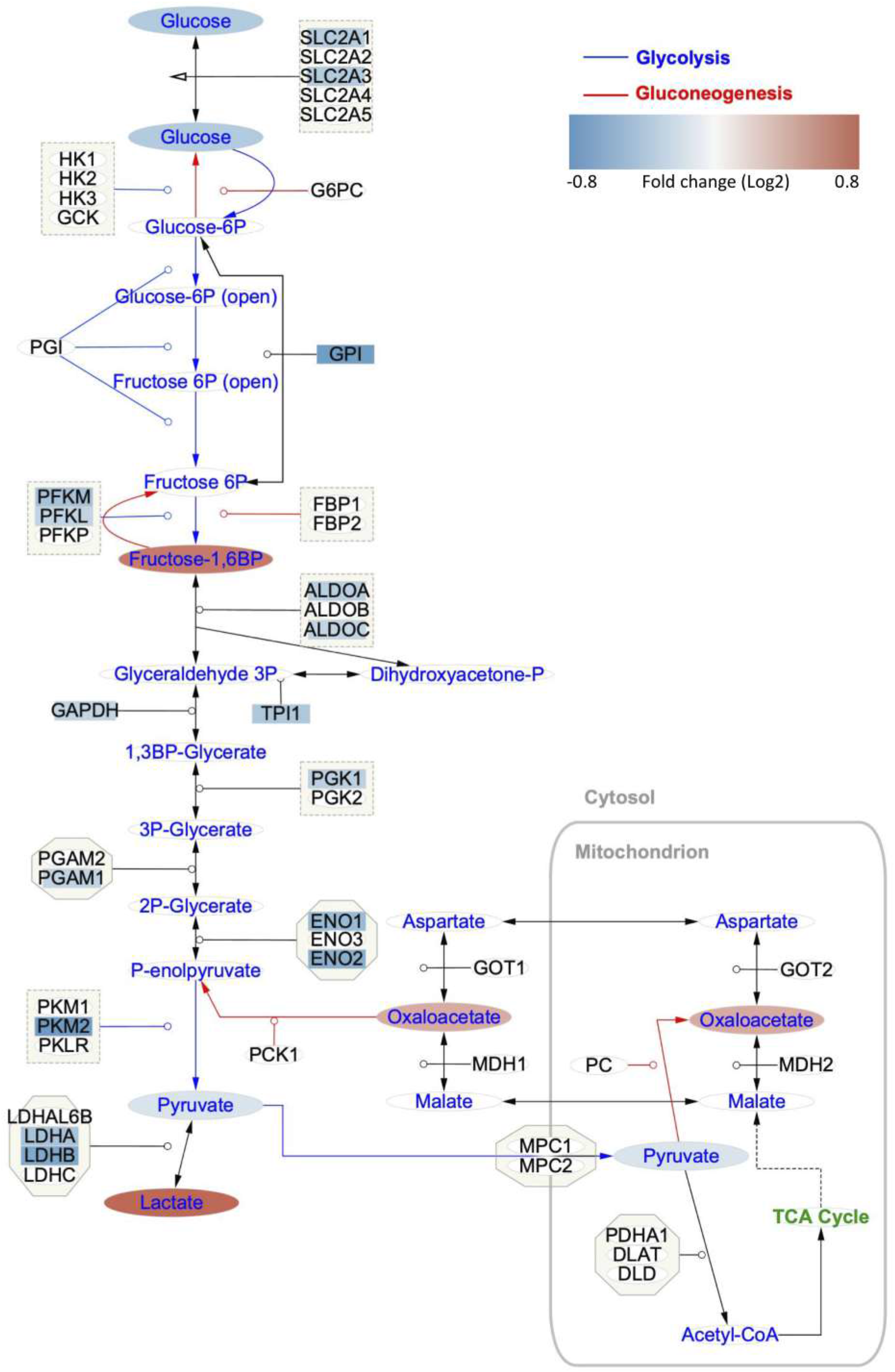
Integrative metabolomic and proteomic characterization of the glycolytic pathway in *PARK2* KO neurons. Glycolysis/gluconeogenesis metabolic pathway created in Cytoscape showing metabolites in circles and proteins in squares. Red and blue colors indicate upregulation and downregulation, respectively. White color indicates metabolites or proteins, which were not detected. The glycolysis pathway is marked by blue arrows and the gluconeogenesis pathway by red colors. Detected proteins: ALDO (aldolase); ENO (enolase); GAPDH (glyceraldehyde 3-phosphate dehydrogenase); LDH (lactate dehydrogenase); PFK (phosphofructokinase); PGAM (phosphoglycerate mutase); PGI/GPI (glucose-6-phosphate isomerase); PGK (phosphoglycerate kinase); PKM (pyruvate kinase isozyme); SLC2A (protein family of glucose transporters); TPI1 (triosephosphate isomerase).

**Figure 7:**
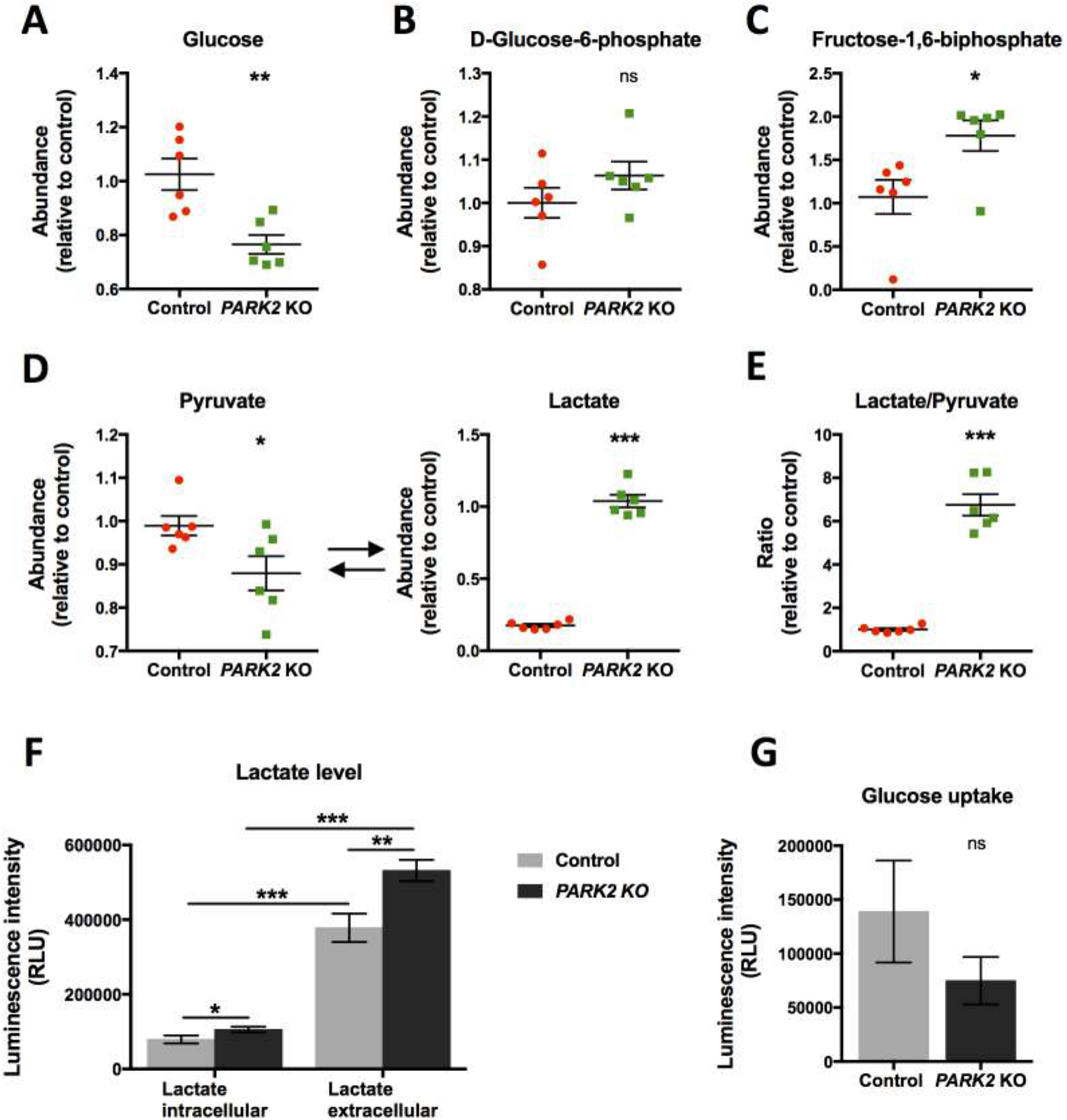
Metabolomic analysis of key intermediates in glycolysis and lactate production. A) Abundance as measured by LC-MS of glucose and further intermediates: B) D-glucose-6-phosphate and C) fructose-1,6-bisphosphate in *PARK2* KO and control neurons. D) Comparison of the levels of pyruvate and lactate and E) ratio between lactate and pyruvate in the *PARK2* KO neurons compared to healthy controls. Levels of F) intracellular and extracellular lactate and G) glucose uptake in *PARK2* KO neurons measured using bioluminescent assays. Data are presented as mean ± SEM, data from 3 independent differentiations, significant differences are indicated by *p < 0.05, **p < 0.01, ***p < 0.001, ns: not significant, Student’s t-test (A-E and G) or two-way ANOVA followed by Tukey’s post hoc test for multiple comparisons (F).

### Accumulation of short- and long-chain carnitines in *PARK2* KO neurons

To further investigate changes in energy-related metabolic pathways, we examined carnitine homeostasis. The carnitine shuttle (*Fig. 8A*) represents a mechanism by which long-chain fatty acids, impermeable to the mitochondrial membranes, are transported into the mitochondrial matrix for β-oxidation and energy production (41). The level of free L-carnitine, which is required for mitochondrial entry of long-chain fatty acids, was significantly reduced in the *PARK2* KO neurons compared with isogenic controls (*Fig. 8B*). In contrast, there was a marked elevation in short- and long-chain acylcarnitines (*Figs. 8C, D*). More specifically, we found an increase in C3-carnitine (propionyl-L-carnitine) and C4-carnitine (O-butanoyl-L-carnitine) as well as unsaturated fatty acid carnitine conjugate profile, palmitoyl-L-carnitine (C16), suggesting decreased lipid β-oxidation in *PARK2*-deficient neurons.

**Figure 8:**
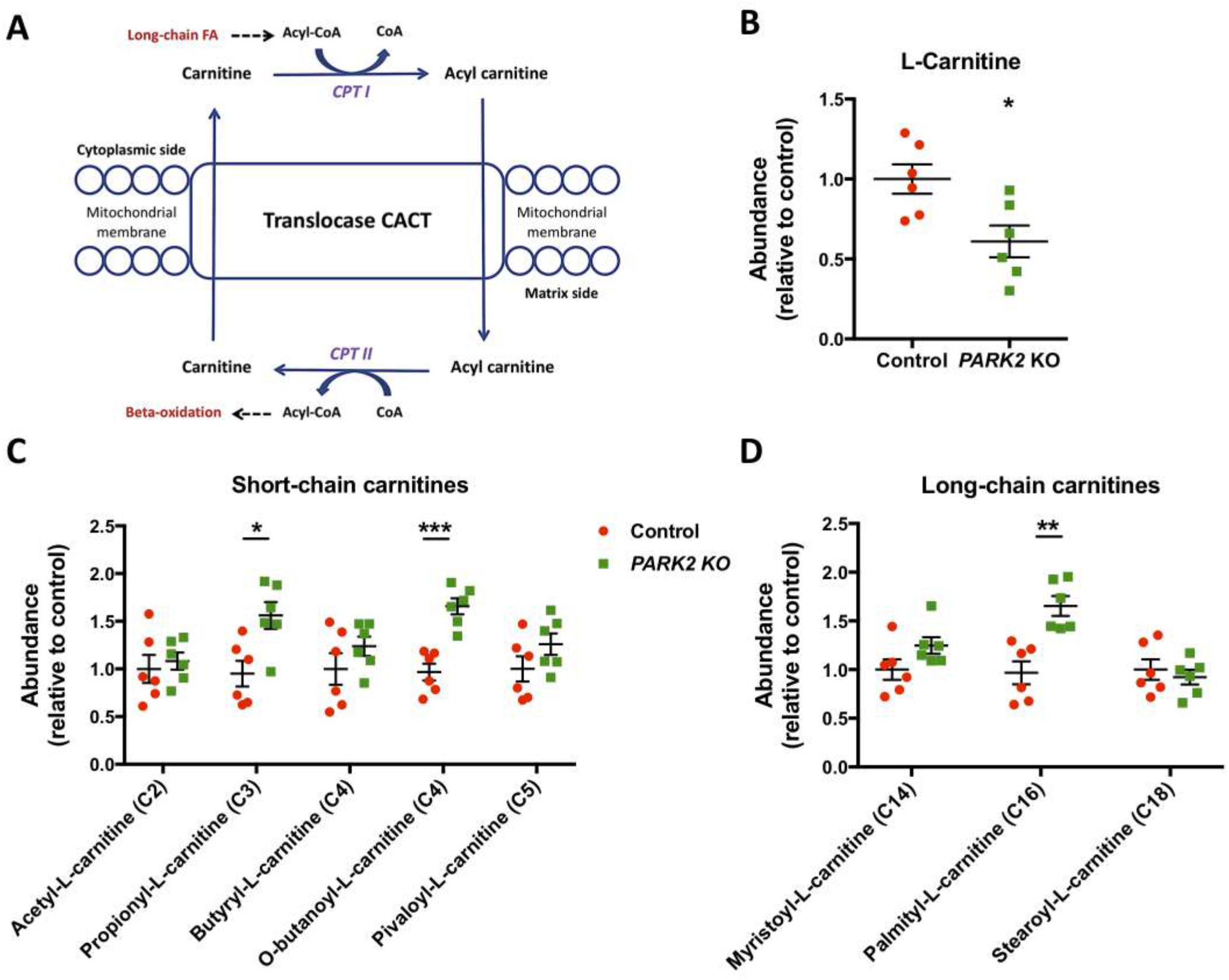
Accumulation of short- and long-chain carnitines in *PARK2* KO neurons. A) Schematic representation of the carnitine shuttle. Levels of B) free L-carnitine, C) short- and D) long-chain acylcarnitines in the *PARK2* KO and control neurons as measured by LC-MS. Data are presented as mean ± SEM, data from 3 independent differentiations, significant differences are indicated by *p < 0.05, **p < 0.01, and ***p < 0.001, Student’s t-test.

### *PARK2* KO neurons display differential oxidative stress-associated metabolic parameters

LC-MS metabolomic profiling suggested that oxidative stress may also be linked to neuronal complications associated with PD. Although the levels of reduced glutathione (GSH) were not significantly changed in the *PARK2* KO neurons (*Fig. 9A*), the level of oxidized glutathione (GSSG) was increased (*Fig. 9B*). Accordingly, *PARK2* KO neurons showed a marked decrease in the GSH/GSSG ratio (*Fig. 9C*), suggesting elevated oxidative stress. In addition, the NADH/NAD ratio, which plays a role in preventing cellular oxidative damage, was imbalanced in the *PARK2* KO neurons (*Fig. 9D*); this was verified with a bioluminescence assay (*Fig. 9E*). In line with the present findings, our previously reported proteomic analysis identified decreased levels of proteins related to superoxide radical degradation and metabolism of ROS, including superoxide dismutase (SOD) 1, catalase, and DJ-1, amongst others (*Fig. 9F*) (26). As imbalance in energy metabolism combined with defective oxidative stress defense is associated with increased ROS production (6, 42, 43), we evaluated ROS levels in the *PARK2* KO neurons and healthy controls. Basal cytosolic ROS production was significantly higher in the *PARK2* KO neurons than in the control cells (*Fig. 9G*), which was also reported in our recent study (27). Overall, we found that oxidative stress was aggravated in the *PARK2* KO neurons.

**Figure 9:**
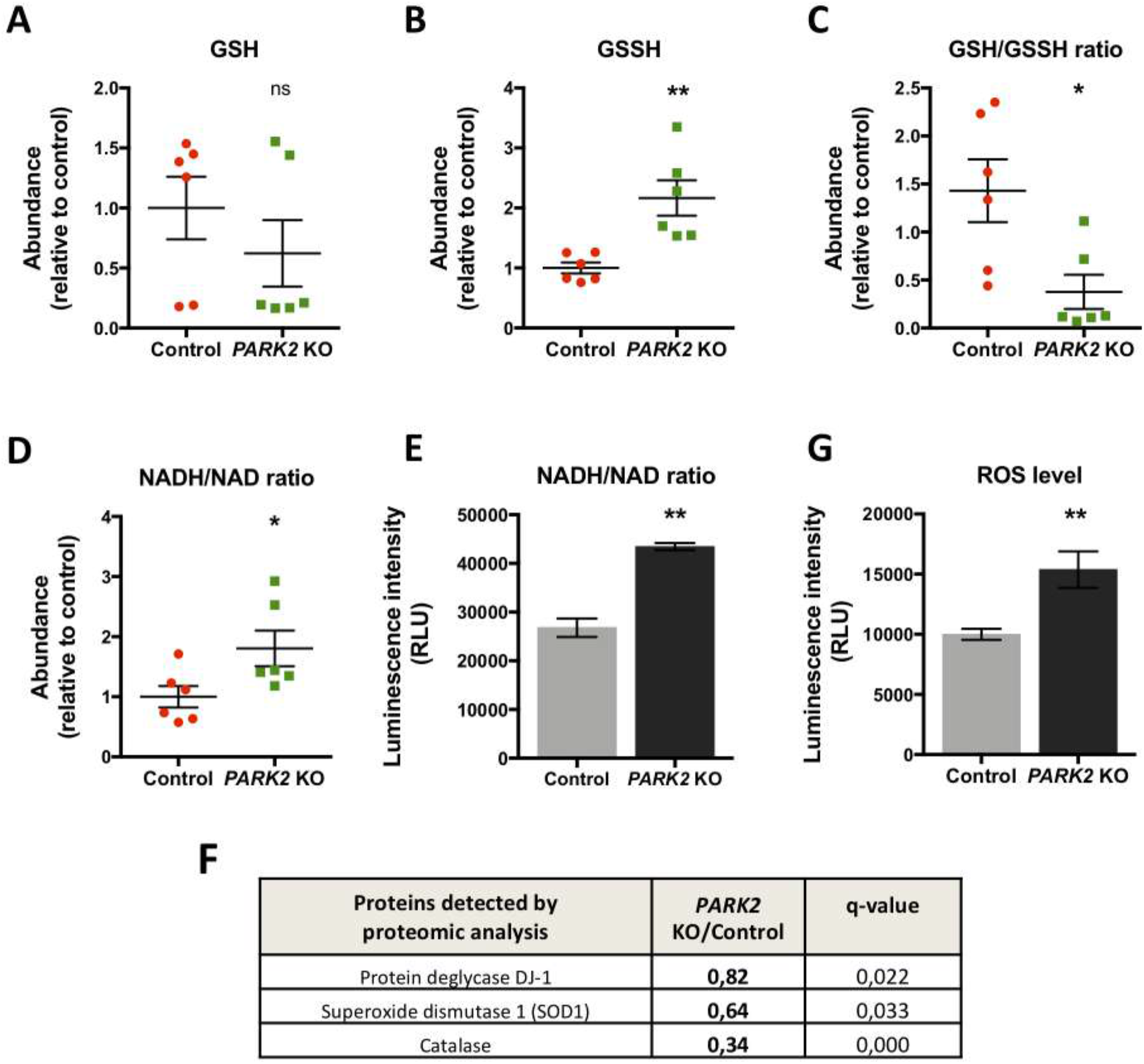
Increased oxidative stress and decreased oxidative stress defense in *PARK2* KO neurons. Levels of A) reduced glutathione (GSH), B) oxidized glutathione (GSSG), and C) the GSH/GSSG ratio revealed by metabolomic analysis. D) NADH/NAD ratio measured by metabolomics and E) verified using a bioluminescence assay. F) Table listing the earlier identified proteins related to oxidative stress defense with the ratio of proteins levels in *PARK2* KO neurons compared to controls (26). G) Levels of cellular ROS production. Data are presented as mean ± SEM, data from 3 independent differentiations, significant differences are indicated by *p < 0.05, **p < 0.01, ns: not significant, Student’s t-test.

## Discussion

Metabolomic analysis is a powerful tool for investigating global changes in the cellular metabolism. Several metabolomic studies have been performed on plasma, cerebrospinal fluid, or blood samples from PD patients with the aim of identifying potential biomarkers (44, 45). To our knowledge, no data has previously been generated on the metabolomic profile of human iPSC-derived neurons with parkin deficiency. We report here that loss of parkin function induces metabolic dysregulation. We show that *PARK2* mutation in human iPSC-derived neurons affected their general metabolome, likely caused by altered mitochondrial and energy homeostasis combined with increased oxidative stress and decreased anti-oxidative response. Energy metabolism in the central nervous system is anticipated to be essential for neuronal health; therefore, impaired energy metabolism plays an important role in the pathogenesis of PD (46). Most of the cellular energy in healthy cells is produced by OXPHOS from the energy precursors produced in the TCA cycle in mitochondria (42, 47). Interestingly, our results showed accumulation of metabolites associated with the TCA cycle in the *PARK2* KO neurons. As the rate of the TCA cycle is regulated primarily by the concentration of ATP and NADH (48), this accumulation of TCA intermediates in *PARK2* KO neurons might be a consequence of the detected low ATP and high NADH levels. Alterations in levels of TCA cycle metabolites have been reported previously in relation to PD. For instance, an increased malate concentration was found in plasma (49), and an increased citrate concentration in the CSF (50) of PD patients. In contrast, decreased levels of several TCA metabolites (citrate, malate, succinate, and isocitrate) have also been detected in brains from PD patients (51). These inconsistencies may be due to the different types of samples being analyzed (neuronal cell cultures, biofluids, brain tissues) or technical variations in sample matrix, sampling, storage, and measurement methods. Moreover, similarly to our results, an increase in concentrations of TCA intermediates was reported in samples from patients with mitochondrial disorders (52, 53). Even if existing literature is slightly divergent, these data point overall to TCA dysregulation resulting in mitochondrial and energy perturbations involved in PD.

Given the perturbed TCA cycle observed in the *PARK2* KO neurons, we further investigated mitochondrial function and found reduced levels of ATP, a critical parameter of cellular energy status. These data were strengthened by the observation of aberrant mitochondrial cristae ultrastructure and reduced MMP in the *PARK2* KO neurons. The inner membrane structures are crucial for ATP production due to maintaining MMP that provides the driving force for ATP synthesis (54). A few other studies, including our own, of PD patient iPSC-derived neurons with *PARK2* mutation have also reported aberrant mitochondrial morphology and function (26, 55, 56), and several prior studies have addressed changes in energy metabolism associated with PD. The fall in ATP has been reported in different experimental PD models, such as MPTP mouse model (57) or skin fibroblasts isolated from PD patients (58).

The altered energy and mitochondrial metabolism described in the present study verifies and complements our earlier study in which we reported mitochondria-specific proteomic and phospho-proteomic changes caused by *PARK2* KO. We found several mitochondrial protein changes combined with disturbances in mitochondrial morphology and respiration, among other (26).

Glycolysis is a process in which glucose is converted into pyruvate or lactate depending on whether it occurs under aerobic or anaerobic conditions. The regulation of glycolysis is complex and occurs at multiple stages (59). In the present study we detected dysregulation of glucose metabolism in *PARK2* KO neurons at various levels of the glycolytic pathway. We found a significant decrease in levels of both glucose (the initial substrate of glycolysis) and pyruvate (the end product of aerobic glycolysis), which was not driven by changes in glucose uptake. This together with the reduced levels of enzymes involved in the glycolytic regulation, detected by proteomic analysis, is indicative of a decrease in glycolytic capacity in *PARK2* KO neurons. These results are in line with our previous data showing that *PARK2* KO causes a significant decrease in extracellular acidification rate (glucose as a substrate), which strongly indicates a decrease in anaerobic glycolysis (26). Fructose-1,6-bisphosphate, which level was elevated in *PARK2* KO cells, is one of the most significant regulators of glycolysis. This phosphorylated fructose sugar is an intermediate of the glycolysis pathway and provides feedforward stimulation as an allosteric activator of two key enzymes of glycolysis: phosphofructokinase (PFK) and pyruvate kinase (PK) (60). Increased level of fructose-1,6-bisphosphate may represent a compensatory mechanism since levels of both key enzymes were significantly reduced in the *PARK2* KO neurons. Further, we showed that the ratio between lactate and pyruvate was significantly elevated in *PARK2* KO neurons compared to healthy controls. This can be an indicator of alterations in protein degradation (61) as well as decreased lactate clearance (62). In addition, a buildup of lactic acid can cause suppression of glycolytic enzymes (63), which to some extent may explain their low level in *PARK2* KO neurons. An increase in lactate levels may be a result of mitochondrial dysfunction promoting the conversion of pyruvate to lactate, which has been reported for mice (64). However, our recent paper showed that the basal oxygen consumption rate was significantly reduced in *PARK2* KO neurons respiring on lactate, which is indicative of impaired lactate-supported respiration (26). It is worth considering that the changed lactate-pyruvate ratio found for *PARK2* KO neurons may be caused by buildup of astrocyte-derived lactate. This assumption is meaningful since there is evidence that astrocyte-derived lactate is an important energy source of the human brain that also may serve as a neuroprotective factor in pathological conditions (65, 66). Overall, the present metabolomic findings are consistent with our previously reported proteomic data, suggesting impaired glycolysis and disturbances in lactate metabolism (26). In general, our data show that the combination of proteomics and metabolomics represent a very valuable platform for better understanding of energy metabolism and disease mechanisms.

Carnitine also plays a critical role in energy production due to transporting long-chain fatty acids into the mitochondria where they can be oxidized (67, 68). The reduction in L-carnitine level and the elevation in some short- and long-chain acylcarnitines in the *PARK2* KO neurons reflect an underling disturbance of mitochondrial fatty acid β-oxidation, suggesting an impaired lipid-handling capacity (41, 69). The observed accumulation of short- and long-chain fatty acid carnitine conjugates and decreased L-carnitine also suggest defects in the carnitine shuttle and hence impaired mitochondrial import of fatty acids. These defects may lead to detrimental effects on ATP production as the import of long-chain fatty acids into the mitochondria is rate-limiting for lipid β-oxidation (70). Increased levels of saturated chain acylcarnitines have been reported in patients with metabolic disorders (70) although another study showed the opposite – a reduction in acetylcarnitines (71). Further studies are warranted in this matter due to the conflicting literature, but all these results demonstrate energy dysregulation of fatty acid oxidation.

Oxidative stress is a condition associated with an increased rate of cellular damage and is manifested by the excessive production of ROS with insufficient or defective antioxidant defense systems (72). Oxidative stress causes profound alterations in various biological structures, including cellular membranes, lipids, proteins, and nucleic acids (73). We have found substantially decreased levels of several antioxidant enzymes (26) as well as increased mitochondrial ROS levels in the *PARK2* KO neurons. These results are in accordance with existing literature indicating generally increased oxidative stress in the peripheral blood of PD patients (74). GSH is a vital cellular protective antioxidant, and the cellular ratio of GSH to GSSG is often used as a marker of cellular toxicity (75). The reduced glutathione redox ratio (GSH/GSSG) in *PARK2* KO neurons is consistent with the concept of oxidative stress being a major component of the pathogenesis of nigral cell death in PD (6, 76). NADH and NAD are fundamental molecules in metabolism and redox signaling (74). We found perturbed balance between NADH and NAD molecules, pointing to a significantly higher level of NADH than NAD in the *PARK2* KO neurons. The consequence of NADH/NAD redox imbalance is initially reductive stress, which eventually leads to oxidative stress and oxidative damage to macromolecules such as DNA, lipids, and proteins (77). In line with our findings, loss of redox homeostasis has previously been associated with many pathological conditions, including PD (78).

In conclusion, in this study we sought to obtain a comprehensive description of the metabolomic changes in a *PARK2* KO iPSC model of PD. Taken together, our results suggest that mitochondrial defects and subsequent changes in energy metabolism and oxidative stress combined with inefficient oxidative stress defense mechanisms may be associated with cellular dysregulation in PD and could represent a potential therapeutic target.

## Acknowledgements

The authors would like to thank Dorte Lyholmer, Nadine Becker-von Buch, Ulla Melchior Hansen, and Maria Pihl for excellent technical assistance, and Dr. Claire Gudex for editing the manuscript.

The research leading to these results was supported by the Innovation Fund Denmark (BrainStem; 4108-00008A), H. Lundbeck A/S, the Danish Parkinson Foundation, the Jascha Foundation, the A.P. Møller Foundation for the Advancement of Medical Science (15-396, 14-427), and the Faculty of Health Sciences at the University of Southern Denmark.

The live imaging experiments reported in this paper were performed at DaMBIC, a bioimaging research core facility at the University of Southern Denmark. DaMBIC was established by an equipment grant from the Danish Agency for Science Technology and Innovation and by internal funding from the University of Southern Denmark.

## Methods

### Ethics statement

The Research Ethics Committee of the Region of Southern Denmark approved the study prior to initiation (S-20130101). All use of human stem cells was performed in accordance with the Danish national regulations, the ethical guidelines issued by the Network of European CNS Transplantation and Restoration (NECTAR) and the International Society for Stem Cell Research (ISSCR).

### *In vitro* propagation and differentiation of neural stem cells (NSCs)

*PARK2* KO and healthy isogenic control NSC cell lines were provided by XCell Science Inc. (Novato, CA, USA) (79). NSC lines were propagated according to well-established, standard protocols using Geltrex-(Thermo Fisher)-coated plates in Neurobasal Medium (Thermo Fisher) supplemented with NEAA, GlutaMax-I, B27, supplement (Thermo Fisher), penicilin-streptomycin, and bFGF. Cells were enzymatically passaged with Accutase (Thermo Fisher) when 80-90% confluent. NSCs were differentiated according to a commercially available dopaminergic differentiation kit (XCell Science Inc., Novato, CA, USA) for at least 25 days. Differentiation was divided into two parts: an induction phase, where NSCs were differentiated into dopaminergic precursors, and a maturation phase, where the dopaminergic precursor cells were differentiated into mature dopaminergic neurons. The differentiations were carried out at 37°C in a low O_2_ environment (5% CO_2_, 92% N2, and 3% O_2_). The cells were seeded onto wells coated with poly-L-ornithine (Sigma) and laminin (Thermo Fisher) at a density of 50,000 cells/cm^2^. Complete DOPA Induction Medium (XCell Science) supplemented with 200 ng/ml human recombinant Sonic Hedgehog (Peprotech) was changed every second day for the first nine days of differentiation. The cells were passaged at day 5 and day 10 and seeded at a desired cell density. The medium was switched to Complete DOPA Maturation Medium A and B (XCell Science) at day 10 and 16, respectively.

### Mass spectrometry (MS)-based metabolomics

#### Sample collection and metabolite extraction

Cells were differentiated according to the DOPA XCell Science protocol in a 6-well culture plate. Differentiated for 25 days, neurons from three independent differentiations in technical duplicates were collected by extraction in a polar, ice-cold solvent (50% methanol (Sigma), 30% acetonitrile (Sigma), 20% water (Merck Millipore), and 50 pmol/ml of amino acid internal standard MIX AA (Cambridge). The wells without cells were included as signal control samples. Briefly, after removing media from wells, the cells were washed 3x in ice-cold PBS (ThermoFisher). Then, 1 ml of ice-cold solvent with internal standard MIX AA was added to each well, and the cell metabolites were extracted from the samples by placing the cell culture plate on a rocking shaker on ice for 5 min. The cell extract was transferred to safe-lock Eppendorf tubes and centrifuged at 10,000 rpm, 4°C for 15 min. Supernatants were transferred to new safe-lock Eppendorf tubes and stored at −20°C until further analysis.

#### Liquid chromatography-mass spectrometry (LC-MS) analysis and data extraction

Samples were resuspended in 20 μl 1% formic acid (FA) in water before centrifugation at 16,000 g for 5 min and transfer of 15 μl to 2 ml HPLC vial containing 100 μl inserts. The samples were analyzed on LC-MS using two different columns; an Agilent Zorbax Eclipse Plus C18 column (2.1 × 150 mm, 1.8 μm) with a 5 mm guard and a Discovery HSF5 HPLC Column (2.1 × 150 mm, 3 μm) with a 20 mm guard. 1 μl was injected on a Agilent 1290 Ininity HPLC system (Agilent Technologies, Santa Clara, CA) using the C18 column, which was kept at 40°C. A flow rate of 300 μl/min was delivered to the column with the following solvent composition: A (0.1 % FA, water) and B (0.1% FA, acetonitrile): 0-2 min (97% A), 2-8 min (97-60% A), 8-12 min (60-10% A), 12-14.5 min (10% A), 14.5-15 min (10-97% A) and 15-18 min (97% A). The HSF5 column was kept at 40°C and was used to inject 0.5 μl. With this setup, a flow rate of 300 μl/min was used with the following solvent composition: A (0.1 % FA, water) and B (0.1% FA, acetonitrile): 0-3 min (100% A), 3-15 min (100-60% A), 15-16 min (60-30% A), 16-17 min (30% A), 17-18 min (30-100% A), and 18-23% (100% A). The flow was coupled to a 6530B quadrupole time of flight mass spectrometer (Agilent Technologies) operated in both negative and positive ion mode (only using C18 column setup) and scanning from 70-1050 m/z with the following settings: 3 scans/sec, gas temp at 325°C, drying gas at 8 l/min, nebulizer at 35 psig, sheath gas temp at 350°C, sheath gas flow at 11 min/l, VCap at 3500 V, fragmentor at 125 V, and skimmer at 65 V. Each spectrum was internally calibrated during analysis using the signals of purine and Hexakis 1H,1H,3H-tetrafluoropropoxy phosphazine, which was delivered to a second needle in the ion source by an isocratic pump running with a flow of 20 μl/min. A pooled sample containing equal volume from each sample was acquired in “all-ion” mode using 0, 10, 20, and 40 V in collision energy. Using this data file, the compounds were putatively annotated to MSI level 3 (80) in Masshunter Qualitative Analysis B.08.00 (81). The resulting list of compounds was then imported to MzMine for annotation. The raw data were converted to mzXML format using MSConvert and processed using MzMine 2 with the chromatogram deconvolution module Wavelets ADAP (82).

#### General data filtering and processing

Compounds with an average abundance of less than 5 times as compared with the signal control extraction samples, were excluded. Compound lists were uploaded to MetaboAnalyst.ca (83). The signals from each analytical setup were normalized by using signal sum. Compound lists from the different setups were merged prior to log2 transformation and auto scaling in MetaboAnalyst software.

The Cytoscape StringApp was used to visualize the differentially expressed proteins and metabolites of the glycolytic pathway, which was imported from WikiPathways (84).

### MMP evaluation

Mitochondrial membrane potential (MMP) was investigated using the TMRE Mitochondrial Membrane Potential Assay Kit (Abcam) according to the manufacturer’s instructions. Briefly, cells were plated parallel in a 24-well culture plate with glass coverslips (for live imaging staining) and in a 96-well culture plate (for microplate reader analysis). TMRE solution was added to cells, and they were incubated for 20 min at 37°C. The staining at 24-well plates was analyzed at Ex/Em = 549/575 nm by confocal microscopy using a FluoView FV1000MPE – multiphoton laser confocal microscope (Olympus) with 60X magnification. The staining at 24-well plates was subsequently analyzed at Ex/Em = 549/575 nm by microplate spectrophotometry (SoftMax1 Pro software; Molecular Devices).

### ATP evaluation

Intracellular ATP level was measured using ATPlite Assay Kit (Perkin Elmer) according to the manufacturer’s instructions. Briefly, cells were plated in a 96-well plate (40,000 cells/well) and were lysed using mammalian lysis cell buffer (Perkin Elmer) and then shaken for 5 min in an orbital shaker at 700 rpm. Substrate solution was added to the wells and the cells were again shaken for 5 min in an orbital shaker at 700 rpm. The plate was dark-adapted for 10 min, and the luminescence signal was recorded using an Orion L Microplate Luminometer (Titertek Berthold).

### Lactate evaluation

Intracellular and extracellular L-lactate levels were measured using Lactate-Glo™ Assay Assay Kit (Promega) according to the manufacturer’s instructions. Briefly, cells were plated in a 96-well plate (40,000 cells/well). Lactate Detection Reagent was added to each well and the plate was briefly shaken for 60 sec. The plate was incubated for 60 min at RT and the luminescence signal was recorded using an Orion L Microplate Luminometer (Titertek Berthold).

### Glucose uptake evaluation

Glucose uptake was assessed using Glucose Uptake-Glo™ Assay Kit (Promega) according to the manufacturer’s instructions. Briefly, cells were plated in a 96-well plate (40,000 cells/well). The cells were washed with PBS, and then 1mM 2DG was added to each well. The plate was briefly shaken and then incubated for 10 min at room temperature (RT). Stop Buffer was added, and the plate was briefly shaken. Neutralization Buffer and 2DG6P Detection Reagent were then added to each well, and the plate was briefly shaken. The plate was incubated for 2 h at RT, and the luminescence signal was recorded using an Orion L Microplate Luminometer (Titertek Berthold).

### NAD and NADH evaluation

NAD and NADH levels were measured using NAD/NADH-Glo™ Assay Kit (Promega) according to the manufacturer’s instructions. Briefly, cells were plated in a 96-well plate (20,000 cells/well). NAD/NADH-Glo™ Detection Reagent was added to each well, and the plate was briefly shaken. The plate was incubated for 1 h at RT, and the luminescence signal was recorded using an Orion L Microplate Luminometer (Titertek Berthold).

### ROS evaluation

ROS level was measured using ROS-Glo™ Assay Kit (Promega) according to the manufacturer’s instructions. Briefly, cells were plated in a 96-well plate (5,000 cells/well). Media were transferred to a new 96-well plate, and H_2_O_2_ substrate was added to each media sample and incubated for 1 h at 37°C. ROS-Glo™ Detection Solution was then added, the plate was incubated for 20 min at RT, and the luminescence signal was recorded using an Orion L Microplate Luminometer (Titertek Berthold).

### Sample preparation for transmission electron microscopy (TEM)

Cells seeded on 13 mm Thermanox plastic coverslips (Nunc) were primarily fixed in 3% glutaraldehyde (Merck) in 0.1 M sodium phosphate buffer with pH 7.2 at 4°C for 1 h and stored in a 0.1 M Na-phosphate buffer at 4°C until further analysis. When ready, the cells were embedded in 4% agar at 45°C (Sigma) under the stereomicroscope and cut into 1-2 mm^3^ blocks, which were then washed with 0.1 M Na-phosphate buffer followed by post-fixation in 1% osmium tetroxide in 0.1 M Na-phosphate buffer (pH 7.2) for 1 h at RT. Cells were washed in MilliQ water, followed by stepwise dehydration in a series of ascending ethanol concentrations ranging from 50-99% EtOH. Propylene oxide (Merck) was then used as an intermediate to allow infiltration with Epon (812 Resin, TAAB). The following day, the agar blocks were placed in flat molds in pure Epon, cured at 60°C for 24 h. Approximately eight semi-thin sections (2 μm) from one block were cut on an ultramicrotome with a glass knife (Leica, Reichert Ultracut UTC). These were stained with 1% toluidine blue in 1% Borax and evaluated by light microscopy to locate areas with adequate number of cells for further processing. Ultrathin sections (70 nm) were cut on the ultramicrotome with a diamond knife (Jumdi, 2 mm) and then collected onto TEM copper grids (Gilder) and stained with 2% uranyl acetate (Polyscience) and 1% lead citrate (Reynolds 1963). The samples were evaluated, and the images were collected using a Philips CM100 transmission electron microscope equipped with a Morada digital camera equipment and iTEM software system.

### Morphometric analysis of TEM images

Six TEM grids from each cell line were used for analysis, and ten spots were randomly chosen at low magnification. Images of each spot were captured using high magnification (19,000X). To estimate mitochondrial abundance, the organelles were counted manually on each micrograph and normalized to cytoplasm area. 30 images were used for each cell line. For further measurement of cristae perimeter/external perimeter ratio, individual organelles were selected and processed manually in ImageJ software using a freehand selection tool, which allows creating irregularly shaped contour. In the total 35-40 organelles from each cell line were analyzed.

### Statistical analysis

Statistical analysis was performed using GraphPad Prism 7.0 software. To analyze the data, two-tailed unpaired Student’s t-test and one- or two-way ANOVA with multiple comparison test were applied. Data were considered statistically significant at p < 0.05 (*), p < 0.01 (**), and p < 0.001 (***). Data are presented as mean ± standard error of the mean (SEM). Metabolomic fold change data were analyzed in MetaboAnalyst software using Student’s t-test.

### Availability of data and material

The remaining datasets used and/or analyzed during the current study are available from the corresponding author on request.

---

**Supplementary Table 1: Changes in metabolites identified by LC-MS-based metabolomic analysis.**

Fold changes (FC) of all detected metabolites levels in *PARK2* KO neurons compared to that of controls. Red and blue colors indicate upregulation and downregulation, respectively. q-value (FDR-adjusted p-value), log2 transformation, and auto scaling; data based on 3 independent experiments.

