## Supplementary Table 1 for "Identification of bioactive metabolites in human iPSC-derived dopaminergic neurons with PARK2 mutation: altered mitochondrial and energy metabolism"

**Supplementary Table 1:** All detected metabolites

| Metabolite | FC *PARK2* KO/Control | Log2(FC) | q-value (Student's t-test) | -Log10(q) |
|---|---|---|---|---|
| Pentadecanedioic acid | 0,09674 | -3,36970 | **1,3E-06** | 5,87330 |
| Tetradecanedioic acid | 0,10972 | -3,18810 | **2,1E-06** | 5,65770 |
| Kynurenine | 11,18400 | 3,48330 | **3,8E-06** | 5,41940 |
| Hexadecanedioic acid | 0,15553 | -2,68470 | **3,8E-06** | 5,41940 |
| Decanedioic acid | 0,25337 | -1,98070 | **3,8E-06** | 5,41940 |
| Undecanedioic acid | 0,17345 | -2,52740 | **4,6E-06** | 5,33380 |
| Octadecanedioic acid | 0,23103 | -2,11380 | **0,00007** | 4,14050 |
| Inosine | 1,89210 | 0,91998 | **0,00022** | 3,65170 |
| L-Aspartic acid | 0,67469 | -0,56771 | **0,00022** | 3,65170 |
| Gluconic acid | 4,39980 | 2,13740 | **0,00024** | 3,61200 |
| L-Lactic acid (Lactate) | 1,68950 | 0,75657 | **0,00028** | 3,55330 |
| Oxoglutaric acid (α-Ketoglutarate ) | 2,95520 | 1,56330 | **0,00137** | 2,86330 |
| Succinic acid (Succinate) | 2,55920 | 1,35570 | **0,00184** | 2,73610 |
| L-Glutamic acid | 1,64640 | 0,71930 | **0,00184** | 2,73610 |
| Fumaric acid (Fumarate) | 1,87240 | 0,90491 | **0,00214** | 2,66890 |
| Hypoxanthine | 1,58940 | 0,66848 | **0,00244** | 2,61260 |
| D-4'-Phosphopantothenate | 0,56992 | -0,81118 | **0,00294** | 2,53140 |
| Sphingosine | 1,75960 | 0,81521 | **0,00532** | 2,27440 |
| Citric acid (Citrate) | 1,31760 | 0,39788 | **0,00572** | 2,24280 |
| O-butanoyl-carnitine (C4) | 1,65640 | 0,72803 | **0,00662** | 2,17920 |
| N-Acetyl-L-Histidine | 0,79045 | -0,33925 | **0,00678** | 2,16900 |
| L-Malic acid (Malate) | 1,59720 | 0,67555 | **0,00905** | 2,04340 |
| Adenosine | 2,66150 | 1,41220 | **0,01091** | 1,96240 |
| LysoPE (0:0/20:0) | 0,56810 | -0,81578 | **0,01380** | 1,86010 |
| LysoPC (18:3) | 0,59753 | -0,74291 | **0,01380** | 1,86010 |
| Oxidized Glutathione (GSSH) | 1,46700 | 0,55291 | **0,01380** | 1,86010 |
| Isocitrate | 1,67600 | 0,74506 | **0,01826** | 1,73840 |
| Palmityl-L-carnitine (C16) | 1,65330 | 0,72536 | **0,01826** | 1,73840 |
| LysoPC (18:0) | 0,56952 | -0,81219 | **0,01956** | 1,70870 |
| LysoPC (18:1) | 0,70963 | -0,49486 | **0,02171** | 1,66340 |
| Pantothenic Acid | 0,84758 | -0,23858 | **0,02315** | 1,63540 |
| N-Acetylaspartylglutamic acid | 0,82469 | -0,27807 | **0,02348** | 1,62930 |
| Glucose | 0,78679 | -0,34595 | **0,02420** | 1,61610 |
| Flavin adenine dinucleotide (FAD) | 1,30840 | 0,38784 | **0,02509** | 1,60050 |
| L-Histidine | 0,60874 | -0,71610 | **0,02587** | 1,58710 |
| S-Adenosylhomocysteine | 1,50800 | 0,59263 | **0,03268** | 1,48570 |
| Uridine | 0,64427 | -0,63427 | **0,03284** | 1,48360 |
| LysoPC (15:0) | 0,73390 | -0,44635 | **0,03404** | 1,46800 |
| N-Acetyl-L-methionine | 1,24050 | 0,31093 | **0,03570** | 1,44730 |
| LysoPC (16:0) | 0,74769 | -0,41950 | **0,03729** | 1,42840 |
| L-Formylkynurenine | 1,96720 | 0,97612 | **0,03814** | 1,41870 |
| Pyruvate | 0,90645 | -0,14170 | **0,03814** | 1,41870 |
| L-Ascorbic acid | 1,56480 | 0,64599 | **0,04003** | 1,39760 |
| LysoPE (20:5) | 0,77941 | -0,35955 | **0,04003** | 1,39760 |
| L-Carnitine | 0,60992 | -0,71330 | **0,04090** | 1,38830 |

| | | | | |
|---|---|---|---|---|
| **LysoPC (18:2)** | 0,77952 | -0,35934 | **0,04990** | 1,30190 |
| **D-fructose 1 6-bisphosphate** | 1,57560 | 0,65590 | **0,05038** | 1,29770 |
| **LysoPE (00/20:1)** | 0,73926 | -0,43585 | **0,05038** | 1,29770 |
| **LysoPE (22:5)** | 0,78059 | -0,35736 | **0,05038** | 1,29770 |
| **LysoPE (00/22:5)** | 0,78059 | -0,35736 | **0,05038** | 1,29770 |
| **Propionyl-L-carnitine (C3)** | 1,56110 | 0,64252 | **0,05071** | 1,29490 |
| **LysoPE (18:4)** | 0,72158 | -0,47077 | **0,06092** | 1,21520 |
| **L-Tryptophan** | 1,22720 | 0,29542 | **0,08303** | 1,08080 |
| **L-Arginine** | 0,77685 | -0,36429 | **0,09407** | 1,02650 |
| **LysoPE (00/16:0)** | 0,70628 | -0,50168 | **0,09497** | 1,02240 |
| **Guanosine** | 1,24040 | 0,31082 | **0,10298** | 0,98724 |
| **LysoPC (14:0)** | 0,76589 | -0,38480 | **0,11615** | 0,93500 |
| **Myristoylcarnitine (C14)** | 1,24700 | 0,31849 | **0,11903** | 0,92433 |
| **L-Threonine** | 1,08790 | 0,12160 | **0,12238** | 0,91229 |
| **Glycine** | 1,11550 | 0,15775 | **0,12405** | 0,90639 |
| **N-Acetyl-L-phenylalanine** | 0,84519 | -0,24266 | **0,13432** | 0,87187 |
| **L-Glutamine** | 0,84769 | -0,23840 | **0,14935** | 0,82579 |
| **Oxaloacetate** | 1,30990 | 0,38942 | **0,16218** | 0,79001 |
| **N-Acetylaspartate** | 0,90438 | -0,14500 | **0,16218** | 0,79001 |
| **Pivaloylcarnitine (C5)** | 1,25960 | 0,33297 | **0,20410** | 0,69016 |
| **NADH** | 1,56590 | 0,64702 | **0,20794** | 0,68206 |
| **LysoPC (20:4)** | 0,86614 | -0,20733 | **0,21570** | 0,66615 |
| **L-Phenylalanine** | 1,16780 | 0,22374 | **0,23426** | 0,63030 |
| **Creatine** | 0,80608 | -0,31101 | **0,24241** | 0,61545 |
| **Butyryl-L-carnitine (C4)** | 1,23760 | 0,30755 | **0,25630** | 0,59126 |
| **LysoPE (00/24:6)** | 0,82396 | -0,27936 | **0,25630** | 0,59126 |
| **D-Glucose 6-phosphate** | 1,06300 | 0,08810 | **0,26834** | 0,57131 |
| **LysoPE (00/20:2)** | 0,79398 | -0,33283 | **0,27466** | 0,56121 |
| **Glutathione** | 1,11410 | 0,15585 | **0,29045** | 0,53693 |
| **Nicotinamide adenine dinucleotide (NAD)** | 1,30900 | 0,38841 | **0,30562** | 0,51482 |
| **L-Valine** | 0,86336 | -0,21197 | **0,51186** | 0,29085 |
| **Xanthine** | 1,08150 | 0,11307 | **0,52581** | 0,27917 |
| **LysoPE (18:3)** | 0,90431 | -0,14511 | **0,53159** | 0,27442 |
| **Flavine mononucleotide (FMN)** | 1,08140 | 0,11294 | **0,54340** | 0,26488 |
| **Pyroglutamic acid** | 1,05850 | 0,08200 | **0,55478** | 0,25588 |
| **Cytidine** | 0,79830 | -0,32499 | **0,55944** | 0,25224 |
| **Acetylcarnitine (C2)** | 1,08200 | 0,11373 | **0,57812** | 0,23798 |
| **Stearoylcarnitine (C18)** | 0,92261 | -0,11621 | **0,67228** | 0,17245 |
| **L-Serine** | 1,03260 | 0,04627 | **0,70444** | 0,15216 |
| **L-Isoleucine** | 1,03080 | 0,04376 | **0,80949** | 0,09179 |
| **L-Tyrosine** | 1,02440 | 0,03477 | **0,83511** | 0,07826 |
| **L-Methionine** | 0,95282 | -0,06972 | **0,84615** | 0,07255 |
| **L-Proline** | 0,96848 | -0,04621 | **0,84615** | 0,07255 |
| **Adenosine triphosphate (ATP)** | 1,59510 | 0,67365 | **0,91458** | 0,03878 |
| **L-Leucine** | 1,01550 | 0,02223 | **0,91458** | 0,03878 |
| **LysoPE (18:2)** | 1,00200 | 0,00285 | **0,95063** | 0,02199 |
| **LysoPC (20:3)** | 0,99304 | -0,01008 | **0,98423** | 0,00691 |
